# Development of neonatal connectome dynamics and its prediction for cognitive and language outcomes at age 2

**DOI:** 10.1101/2023.08.07.552267

**Authors:** Yuehua Xu, Xuhong Liao, Tianyuan Lei, Miao Cao, Jianlong Zhao, Jiaying Zhang, Tengda Zhao, Qiongling Li, Tina Jeon, Minhui Ouyang, Lina Chalak, Nancy Rollins, Hao Huang, Yong He

## Abstract

The functional connectome of the human brain comprises time-varying network structure that facilitates efficient inter-module communication and support flexible cognitive functions. However, little is known about how the connectome dynamics of the brain emerges and develops at very early stages of human life and whether this dynamics is predictive of neurocognitive outcomes later in life. Here, we employed resting-state functional MRI data from 39 infants (31 to 42 postmenstrual weeks) and a multilayer network model to characterize the development of connectome dynamics during the third trimester and its critical role in predicting future neurocognitive outcomes at 2 years of age. We observed that the modular architecture of baby functional connectomes spontaneously reconfigures over time, with lower network module switching across time primarily in the primary regions and higher module switching mainly in the association areas. With development, the dynamic switching between the brain modules was significantly decreased, primarily located in the lateral precentral gyrus, medial temporal lobe, and subcortical areas. The clustering analysis further revealed that the primary areas displayed a higher developmental rate than the higher-order systems. Using the support vector regression approach, we found that brain connectome dynamics at birth significantly predicted cognitive and language performance at 2 years of age. Our findings highlight the emergence and spatially inhomogeneous maturation of the neonate connectome dynamics, laying a critical neural foundation for the development of cognitive and language skills later in life.

## Introduction

The third trimester is a critical neurodevelopmental stage for the human brain (Rakic 1972, 1995; Tau and Peterson 2010). During this period, the human brain undergoes explosive growth in both structure and function, laying the foundations for cognitive and behavioral development in later life (Cao et al. 2017a; Gilmore et al. 2018; Ouyang et al. 2019a; Zhao et al. 2019). At the microscopic level, the rapid and abundant neural migration, synaptogenesis, and axon growth foster specific neural circuits (Tau and Peterson 2010; Kostovic et al. 2019) supporting primary sensorimotor functions and higher cognitive skills (Dehaene-Lambertz and Spelke 2015). At the macroscopic level, functional connectome mapping studies based on resting-state fMRI (rs-fMRI) have revealed the remarkable reconfiguration of the inter-regional functional connectivity patterns. Specifically, the community structure, which comprises functionally specific and interacting modules, has been observed in the functional connectome of fetuses in utero (Thomason et al. 2014) and preterm and term infants (Cao et al. 2017b). This community structure is supposed to facilitate efficient functional segregation and integration at low wiring costs (Sporns and Betzel 2016). The primary visual, auditory, and sensorimotor modules show adult-like patterns before birth, while the higher-order default-mode and frontoparietal modules exhibit prolonged development after birth (Fransson et al. 2007; Fransson et al. 2009; Doria et al. 2010; Smyser et al. 2010; Cao et al. 2017b). The development of these modular structure promotes functionally segregated processing of the baby brains and makes the functional connectome towards a more organized pattern (Cao et al. 2017b). These findings provide valuable insights into the emergence and development of brain network modules during the third trimester.

Despite the rich evidence on the prenatal development of functional connectomes, most of the previous connectome research has primarily focused on the static (i.e., time-constant) functional networks, largely ignoring the time-varying dynamic patterns of functional connectome. The human brain is a highly dynamic system in nature. Accumulating evidences indicate that the inter-regional functional coordination at rest spontaneously fluctuate at a time scales of seconds or minutes (Hutchison et al. 2013; Liao et al. 2015; Preti et al. 2017). The modular architecture in adult connectomes dynamically reconfigure over time with a high switching among modules in the lateral frontoparietal regions (Liao et al. 2017; Pedersen et al. 2018; Liu et al. 2020). This connectome dynamics maintains efficient communication between network modules (Zalesky et al. 2014) and facilitates a rapid response to potential or ongoing cognitive demands (Barbey 2018; Khambhati et al. 2018; Uddin 2021). Meanwhile, the connectome dynamics varies across individuals (Liao et al. 2017; Liu et al. 2020) and captures individual differences in behavioral and cognitive performance, such as learning capacity (Bassett et al. 2011), executive function (Braun et al. 2015), and cognitive flexibility (Pedersen et al. 2018). Some recent studies have begun to explore the functional connectome dynamics from birth to 2 years of age (Huang et al. 2020; Wen et al. 2020; Yin et al. 2020). These studies reported that the connectivity variability of higher-order brain systems increases with age while the primary systems decreases with age (Wen et al. 2020). Neural flexibility, in terms of modular switching, also increases with age and neural flexibility at 3 months of age is negatively correlated with cognitive ability at approximately 5 years of age (Yin et al. 2020). However, how the connectome dynamics develops before birth and whether this early development shapes neurocognitive outcomes later in life remain to be elucidated.

To address these issues, we analyzed rs-fMRI data from 39 preterm and full-term infants scan aged from 32 to 42 postmenstrual weeks to examine the functional connectome dynamics during the prenatal stage. We further explored the potential associations between the connectome dynamics at birth and future neurocognitive outcomes at around 2 years of age. Specifically, we detected the time-varying modular architecture for each infant using the multilayer modularity framework, which can incorporate the connectivity information between adjacent time points (Mucha et al. 2010). We further used modular variability (Liao et al. 2017) to quantify how brain nodes switch between modules over time. We hypothesized that the network module dynamics would present distinct developmental changes between primary and higher-order systems during the third trimester and predict the neurocognitive outcomes at age 2 years old.

## Materials and Methods

### Participants

We employed rs-fMRI data from 52 healthy preterm and full-term infants. Neonates were recruited from the Parkland Hospitals and underwent MRI scanning during natural sleep without any sedation. Written informed consent was obtained from parents of each participating infant. This study was approved by the Institutional Review Board of the University of Texas Southwestern Medical Center. Detailed exclusion criteria have been described in our previous works (Cao et al. 2017b; Xu et al. 2019; Ouyang et al. 2020). After excluding 13 infants due to excessive motion artifacts (see “Imaging data preprocessing”), we finally included rs-fMRI data from 39 infants with postmenstrual ages ranging from 31.9 to 41.7 weeks at the time of the scan. The demographic data of the infants is described in Table 1.

**Table 1.** Demographic information and neurocognitive outcomes of participating infants.

| Characteristics | | Mean ± SD (range) |
| --- | --- | --- |
| **Imaging**<br>(N=39) | Age at birth (weeks)<sup>a</sup> | 33.0 ± 4.5 (25.0-40.9) |
| | Age at scan (weeks)<sup>a</sup> | 36.9 ± 2.7 (31.9-41.7) |
|  | Male | 28 (72%)<sup>c</sup> |
| | Time from birth to scan (days) | 27.3 ± 26.8 (0-106) |
| **Bayley III**<br>(N=26) | Age at test (months)<sup>b</sup> | 23.2 ± 1.6 (20.3-26.9) |
| | Cognitive scale | 86.7 ± 8.4 (70-110) |
| | Language scale | 87.5 ± 9.1 (71-112) |
| | Motor scale | 91.1 ± 7.3 (73-100) |

<sup>a</sup> postmenstrual age in weeks; <sup>b</sup> chronological age corrected for prematurity; <sup>c</sup> fraction of infants among the whole population.

### Neurocognitive assessments

In this study, 26 of 52 neonates were assessed with the Bayley Scales of Infant and Toddler Development III (Bayley 2006) at approximately 2 years of age, corrected for prematurity (23.1 ± 1.6, 20.3-26.9 months). The Bayley-III test comprises five scales: cognitive, language (expressive and receptive language), and motor (gross and fine motor) scales for infants, as well as social-emotional and adaptive scales from parents’ interviews. The cognitive scale of the Bayley-III test assesses sensorimotor development, concept formation, memory, simple problem-solving, and reasoning skills (Bayley 2006). The language scale of the Bayley-III test includes two subsets: receptive and expressive communication, which measures the child’s ability to understand and use spoken language to follow instructions, label or recognize objects and people based on spoken descriptions (Bayley 2006). The motor scale from the Bayley-III test evaluates both gross and motor skills, such as visual tracking, reaching objects, and the child’s ability to keep balance and jump (Bayley 2006). The neurocognitive assessments were conducted by a certified neurodevelopmental psychologist. The neurocognitive outcomes of the infants in this study is described in Table 1.

### Imaging data acquisitions

All infants were well-fed and had fallen asleep before scanning. To reduce the sound of the scanner, earplugs, earphones, and extra foam padding were applied to the sleeping infants. Images were acquired using a Philips 3 T Achieva MR scanner with an 8-channel SENSE head coil at the Children’s Medical Center at Dallas. The rs-fMRI scans were obtained using a T2-weighted gradient-echo EPI sequence: repetition time = 1500 ms, echo time = 27 ms, flip angle = 80°, in-plane imaging resolution = 2.4×2.4 mm^2^, in-plane field of view = 168×168 mm^2^, slice thickness = 3 mm with no gap, and slice number = 30. A total of 210 whole-brain EPI volumes were acquired. A T2-weighted structural image was acquired with a turbo spin-echo sequence: repetition time = 3000 ms, echo time = 80 ms, in-plane imaging resolution = 1.5×1.5 mm^2^, in-plane field of view = 168×168 mm^2^, slice thickness = 1.6 mm with no gap, and slice number = 65. The acquired T2-weighted image was zero-filled to a 256×256 image matrix.

### Imaging data preprocessing

The rs-fMRI images were preprocessed using toolboxes of Statistical Parametric Mapping (SPM12, http://www.fil.ion.ucl.ac.uk/spm), GRETNA (Wang et al. 2015), and DPARSFA (Yan and Zang 2010). We first discarded the first 15 volumes to allow for the signal to reach a steady state, retaining 195 time points for each infant. The remaining functional data were corrected for the time delay between slices and head motion between volumes. We calculated the framewise displacement (FD) (Power et al. 2012) to evaluate the head motion for each infant. At this stage, data from 12 infants were excluded due to large head motion with criteria of displacement > 5 mm, rotation > 5°, or mean FD > 1 mm. Next, the individual functional data were spatially normalized to a customized template in two steps. First, the individual functional data were coregistered to their corresponding high-resolution T2-weighted structural images using a linear transformation. Then, the individual T2-weighted images were nonlinearly registered to a 37-week brain template (Serag et al. 2012), which corresponds to the average age for all participants. The customized template was then generated by averaging the resultant normalized T2-weighted structural images of all infants and was used as the template for the second registration of individual T2-weighted images. The aligned functional data were spatially normalized to the customized template by applying the transformation parameters estimated during the second registration of T2-weighted images and were resampled to 3-mm isotropic voxels. In addition, prior templates of the cortex, deep gray matter, white matter, and cerebrospinal fluid tissue templates constructed at 37 weeks (Serag et al. 2012) were also registered to the customized template to generate the corresponding tissue masks. Next, the normalized functional imaging data were smoothed with a Gaussian kernel (full width at half-maximum of 4 mm). We further performed the nuisance regression to reduce the effects of head motion and other non-neural signals, including Friston’s 24 head motion parameters (Friston et al. 1996), white matter and cerebrospinal fluid signals, and the global signals. Of note, spike-based regressors were also included in the nuisance regression procedure (Yan et al. 2013; Power et al. 2014) to better control the potential influence of transient head motion. The regressors were defined as the bad volumes with framewise displacement above 0.5 mm and their adjacent volumes (1 back and 2 forward). Here, one infant was further excluded due to that half of the volumes of this infant were bad volumes. Finally, a temporally bandpass filtered (0.01-0.08 Hz) was applied on the residual time series.

### Construction of the dynamic functional connectome

For each infant, we constructed the dynamic functional connectome as follows. First, we defined network nodes via a customized random parcellation, which was obtained by parcellating the gray matter tissue into 256 regions with uniform sizes (Zalesky et al. 2010). Next, the commonly used sliding-window approach was employed to estimate the dynamic functional connectivity between nodal regions (Hutchison et al. 2013; Lurie et al. 2020). The time-dependent functional correlation matrix was estimated as the Pearson’s correlation between time series of brain nodes within each window. The window length was set as 40 TRs (i.e., 60 s) and shifted with a step of 1 TR (i.e., 1.5 s), resulting in a total number of 156 windows. Then, we removed the negative correlation values due to their ambiguous physiological interpretations (Fox et al. 2009; Murphy and Fox 2017) and generated the dynamic weighted functional network by thresholding each windowed correlation matrix with a fixed density of 15%. We also assessed the potential effects of different sliding window lengths (i.e., 100 s) and various network densities (i.e., 10% and 20%) on the main results (see “Validation analysis”).

### Characterizing the functional module dynamics

To track the time-varying functional modular architecture, we used a multilayer network framework (Mucha et al. 2010) to incorporate the functional connectivity information in different time windows. Under this framework, brain nodes in each window were not only connected with nodes in the same time window but also connected with themselves in the two adjacent time windows. The time-varying functional modular architecture was identified by optimizing the multilayer modularity index (*Q*), which is defined as follows:

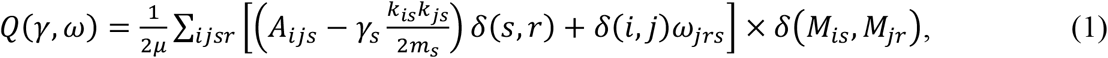

where *i* and *j* represent nodal labels, *s* and *r* represent layer labels. The variable *μ* denotes the total connectivity strength of the multilayer network, including both the intra-layer and inter-layer connectivity strength. For the functional network in layer *s*, the *A*_*ijs*_ represents the functional connectivity strength between node *i* and node *j*, 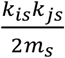 represents the connection probability expected by chance between node *i* and node *j*, and *M*_*is*_ denotes the module assignments of node *i* in the layer *s*. The functional *δ*(*M*_*is*_, *M*_*jr*_) is the Kronecker function that equals 1 if the two variables *M*_*is*_ and *M*_*jr*_ are equal, and equals 0 otherwise. The topological resolution parameter *γ* determines the spatial resolution of the intra-layer module structure, and the temporal coupling parameter *ω* controls the strength of the inter-layer coupling. In our main analysis, the topological resolution parameter *γ* and the temporal coupling parameter ω were set as the default values with *γ = ω* = 1. The choices of other values were also assessed (see “Validation Analysis”). The multilayer community was detected by using the Genlouvain MATLAB package (http://netwiki.amath.unc.edu/GenLouvain).

Then, we employed a measure of modular variability (MV) to quantify how each nodal region switches between modules across time windows (Liao et al. 2017). The larger the modular variability is, the more frequently a brain node switches among modules. For a given node *i*, the modular variability of this node between two windows *k* and *l* was calculated as follows:

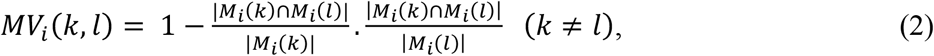

where *T* represents the number of time windows, *M*_*i*_(*k*) and *M*_*i*_(*l*) represent the module affiliations of node *i* in windows *k* and *l*, respectively, |*M*_*i*_(*k*)| denotes the number of nodes included in module *M*_*i*_(*k*), and |*M*_*i*_(*k*) ∩ *M*_*i*_(*l*)| denotes the number of common nodes included in module *M*_*i*_(*k*) and module *M*_*i*_(*l*). We estimated the total modular variability of node *i* across all the time windows (Liao et al. 2017) as:

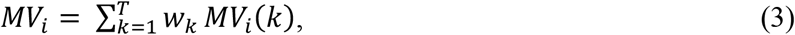

where 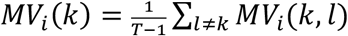 denotes the average modular variability of node *i* between window *k* and all the other windows. Here, we used a normalized weighed coefficient *w*_*k*_ to reduce the bias of potential outlier time windows, which was estimated using adjusted mutual information (Vinh et al. 2010).

Given the heuristic uncertainty of the modularity optimization algorithm (Mucha et al. 2010), we repeated the module detection processes 100 times. The modularity index and modular variability values used for the subsequent analysis were obtained as the averaged values across 100 instances.

### Statistical analysis

To detect age effects on global and nodal dynamic properties, we used the general linear model analysis to quantify the relationship between each dynamic measure and the postmenstrual age as follows:

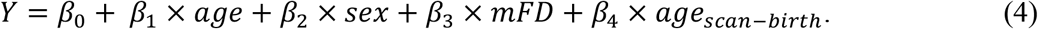

In this model, we also considered three covariates, including sex, mean FD (mFD), and the time interval between birth and the scan. The significant age effect was identified with the significance level of *p* < 0.05. For the nodal analysis, we performed the false discovery rate (FDR) method (Benjamini and Hochberg 1995) to correct for multiple comparisons across nodes.

### Clustering analysis of regional developmental rates of module dynamics

Considering that the development of the module dynamics is spatially heterogeneous, we used a data-driven k-means clustering method (Seber 2009) to classify the network nodes with similar developmental trajectories. For each network node, the age-related beta value that represent the development rate was used as the feature of clustering analysis. The distance between any two brain nodes was defined as the Euclidean distance between their developmental rates. This analysis was repeated with the cluster number varying from 2 to 8, separately. The silhouette value was used to determine the optimal cluster number (Rousseeuw 1987).

### Prediction of neurocognitive outcomes using module dynamics

We further investigated whether the network module dynamics at birth could serve as a biomarker for predicting future neurocognitive outcomes. Briefly, we trained a support vector regression (SVR) model with a linear kernel (Chang and Lin 2011) to separately predict the individual’s cognitive and language scores obtained from the Bayley-III test of each infant at 2 years old. The nodal modular variability at birth were input as features. To evaluate the predictive performance, we employed the 10-fold cross validation (10F-CV) strategy. In the 10F-CV, the data from all infants were divided into 10 subsets.

Specifically, we sorted the infants according to the outcome (i.e., age at scan) and ensures the average age for each subset was nearly same. During each iteration, we used the SVR model derived from the training data (i.e., data from 9 subsets) to predict the Bayley score of the remaining test subset. To assess the prediction accuracy, we calculated partial correlation coefficient between the actual and predicted scores, controlling for age at scan time, sex, mean FD, and the time interval between birth and the scan. To validate that our above split strategy was representative, we tested the prediction accuracy using repeated random 10F-CV. Specifically, the infants were randomly divided into 10 subsets to re-train the SVR model. This procedure was repeated for 1,000 times. The partial correlation values across all 1,000 times were then averaged to represent the overall prediction accuracy. The statistical significance of the prediction was assessed by permutation tests (n = 10, 000). During each permutation instance, we shuffled the Bayley scores across infants before the SVR analysis and re-estimated the partial correlation between the actual and predicted scores. To assess the prediction contribution of all nodes, we re-trained a new SVR model by including all the infants and the contribution weight for each brain node was defined as the absolute value of its weight (Cui and Gong 2018). The SVR model was conducted using the LIBSVM toolbox for MATLAB, with default settings (https://www.csie.ntu.edu.tw/~cjlin/libsvm/).

### Validation analysis

To investigate the reliability of our main results, we evaluated the potential effects of different network analysis strategies, including variations in the sliding window length, network density, and multilayer network parameters. (i) Sliding window length. The optimal selection of sliding window length remains unclear. In the main analysis, we used a recommended length of 60 s to reliably capture the temporal variations in functional networks (Lurie et al. 2020). For validation, we set the window length as 100 s to reconstruct the dynamic network and repeated the analysis. (ii) Network density. We reconstructed the dynamic functional networks with different network densities, including 10% and 20%, separately. (iii) Multilayer network parameters. We evaluated the potential influence by selecting different sets of the temporal parameter *ω* (*ω* = 0.5 and 0.75) and the topological parameter *γ* (*γ* =0.9). When one parameter was reset, the other parameter was retained as the default value (i.e., 1).

## Results

### Module dynamics decreased with development during the prenatal period

At the global level, we found that the modularity index *Q* of the time-varying modular architecture increased significantly with age (Fig. 1A, *t* = 4.97, *p* < 0.001), indicating increasing network segregation during the prenatal period. The global mean values of nodal module variability across the brain decreased significantly with age (Fig. 1A, *t* = -3.19, *p* = 0.003), whereas the corresponding standard deviation significantly increased with age (Fig. 1A, *t* = 3.55, *p* = 0.001), indicating the reduced modular switching and increased spatial heterogeneity with development. At the regional level, we obtained heterogeneous patterns of nodal modular variability by estimating the fitted module variability maps from 32 to 41 postmenstrual weeks (Fig. 1B). Lower modular variability was primarily located in the primary motor areas, and the prefrontal cortex, whereas higher modular variability was mainly located in the lateral and medial frontal and parietal cortices, and the middle temporal gyrus, regardless of age.

**Figure 1.**
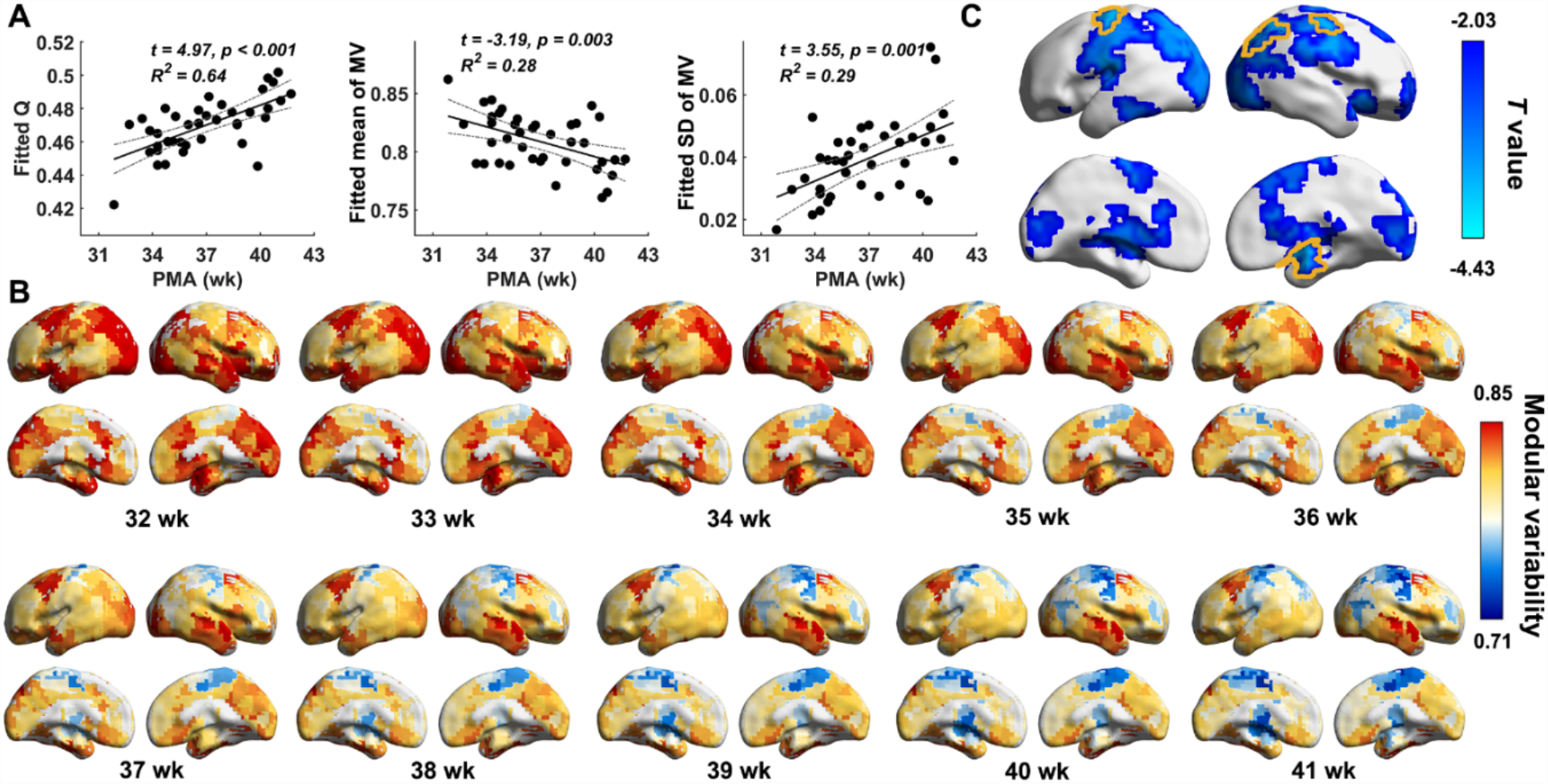
Developmental changes of modular variability during the third trimester. (A) Age-related changes of modularity index *Q*, mean modular variability across the brain, and the corresponding standard deviation. (B) Modular variability maps from 32 to 41 weeks. The maps were estimated as the fitted values from the general linear model, which corrected for influence of sex, head motion parameters (mean FD), and time interval between birth and scan. (C) Age effects on regional modular variability. We only display regions that show significant age-related changes (*p* < 0.05, uncorrected). Yellow curves delineate bran regions that remained significant with correction for multiple comparisons (*p*_*FDR*_ < 0.05). In (B) and (C), results were mapped onto the cortical surface using BrainNet Viewer (Xia et al. 2013). PMA (wk), postmenstrual age in weeks; *Q*, modularity; MV, modular variability; SD, standard deviation; FDR, false discovery rate.

Quantitative analysis revealed significant age-related decreases in modular variability, which were mainly located in the supplementary motor area, precentral gyrus, superior parietal lobe, and medial temporal lobe (Fig. 1C. *p*_*FDR*_ < 0.05, areas delineated with yellow lines, correcting for multiple comparisons).

### Divergent developmental rates between primary and higher-order systems

To delineate the divergent developmental profiles of nodal modular variability across the brain, we employed a k-means clustering method to identify nodal regions with similar developmental curves. Figure 2A shows the inter-regional differences in the developmental rates of modular variability. A two-cluster model was chosen because of its highest silhouette value (Fig. 2B). Cluster 1 primarily contained the sensorimotor areas, lateral occipital, lateral parietal, and subcortical regions (Fig. 2C). Cluster 2 primarily included the lateral and medial frontal regions, medial parietal and occipital regions, and lateral temporal cortex (Fig. 2C). Specifically, Cluster 1 showed a significantly higher developmental rate than Cluster 2 (*t* = -21.96, *p* < 0.001) (Fig. 2D), suggesting the differentiation of the developmental rate between the primary and higher-order systems. For each cluster, we showed the developmental profile of a representative node: the supplementary motor area from Cluster 1 showing significant age-related changes in modular variability (Fig. 2E, *t* = -3.83, *p* < 0.001) and one medial frontal node from Cluster 2 showing non-significant age-related changes in modular variability (Fig. 2E, *t* = -1.15, *p* = 0.257).

**Figure 2.**
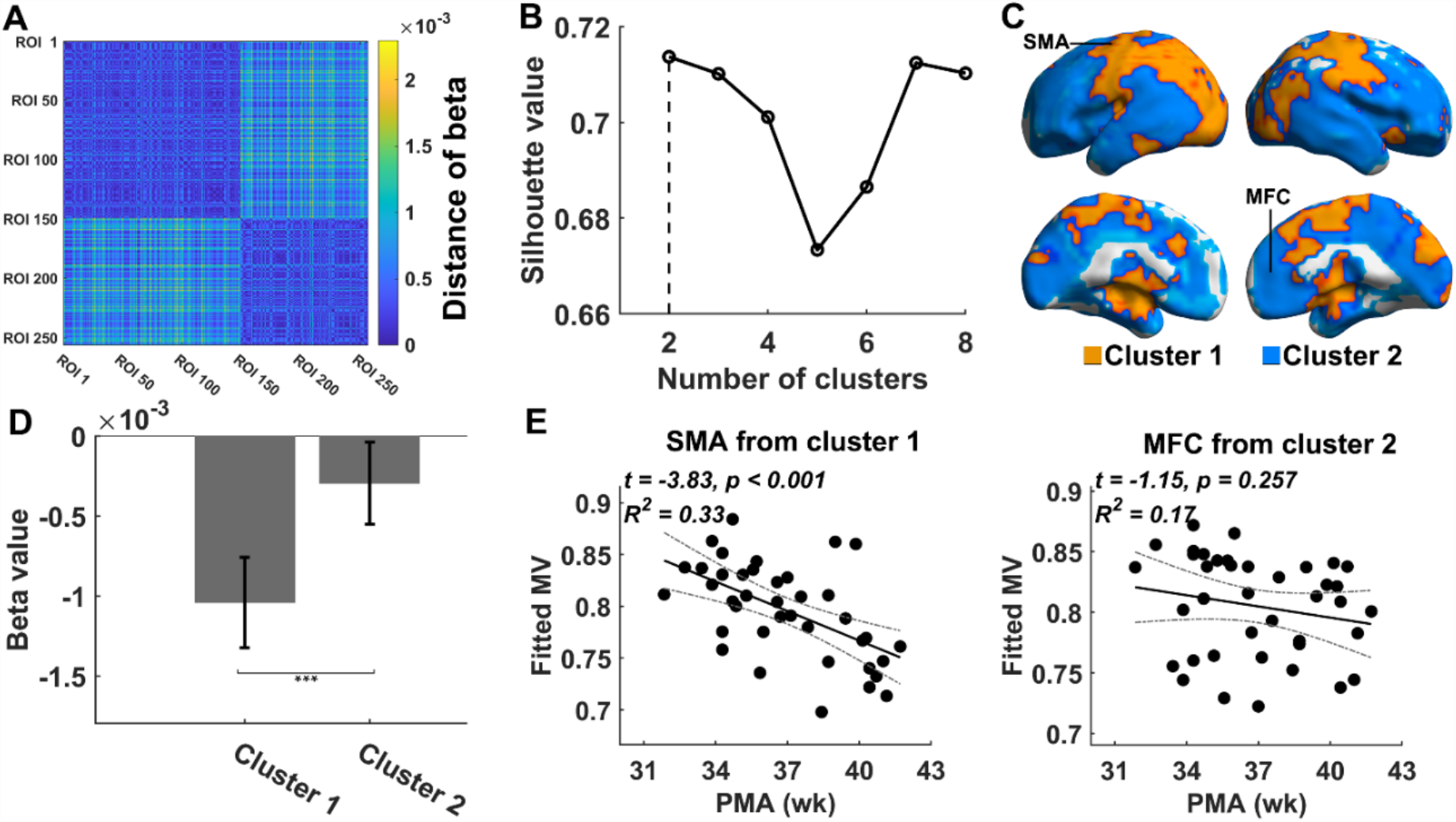
Clustering analysis based on developmental rates of nodal modular variability. (A) Inter-nodal Euclidean distance matrix regarding developmental rates. (B) Silhouette values for the clustering analysis with different clustering numbers. The optimal choice was observed with the 2-cluster model. (C) Spatial locations of two clusters. Two representative nodes are labelled. (D) Different developmental rates of nodal modular variability between two clusters. ***, *p* < 0.001. (E) Age-related changes for two representative nodes. One represents node was selected for each cluster. SMA, supplementary motor area; MFC, medial frontal area; MV, modular variability; PMA (wk), postmenstrual age in weeks.

### Prediction of neurocognitive outcomes using module dynamics

We employed the SVR model with the 10F-CV strategy to evaluate whether the connectome dynamics at birth could serve as a biomarker for individualized prediction of neurocognitive outcomes. We found that the module variability maps at birth significantly predicted both cognitive scores (Fig. 3A, *r* = 0.32, *p*_*perm*_ = 0.021) and language scores (Fig. 3D, *r* = 0.43, *p*_*perm*_ = 0.003) at 2 years old. Repeated random 10F-CV (1,000 runs) demonstrated similar results. (Fig.3B, mean *r* = 0.28, *p*_*perm*_ = 0.044 for cognition prediction, Fig.3E, mean *r* = 0.44, *p*_*perm*_ = 0.006 for language prediction). The brain nodes with high contributions to the cognitive score prediction were mainly located in several default-mode regions (e.g., lateral and medial prefrontal cortex, precuneus, and middle temporal gyrus), lateral and medial occipital cortex, and insula (Fig. 3C). The brain nodes with high contributions to the language score prediction were mainly located in the inferior frontal gyrus, supramarginal gyrus, superior temporal gyrus, lateral and medial occipital cortex, and insula (Fig. 3F). However, the motor score was not able to be predicted from the modular variability (*r* = 0.08, *p*_*perm*_ = 0.269).

**Figure 3.**
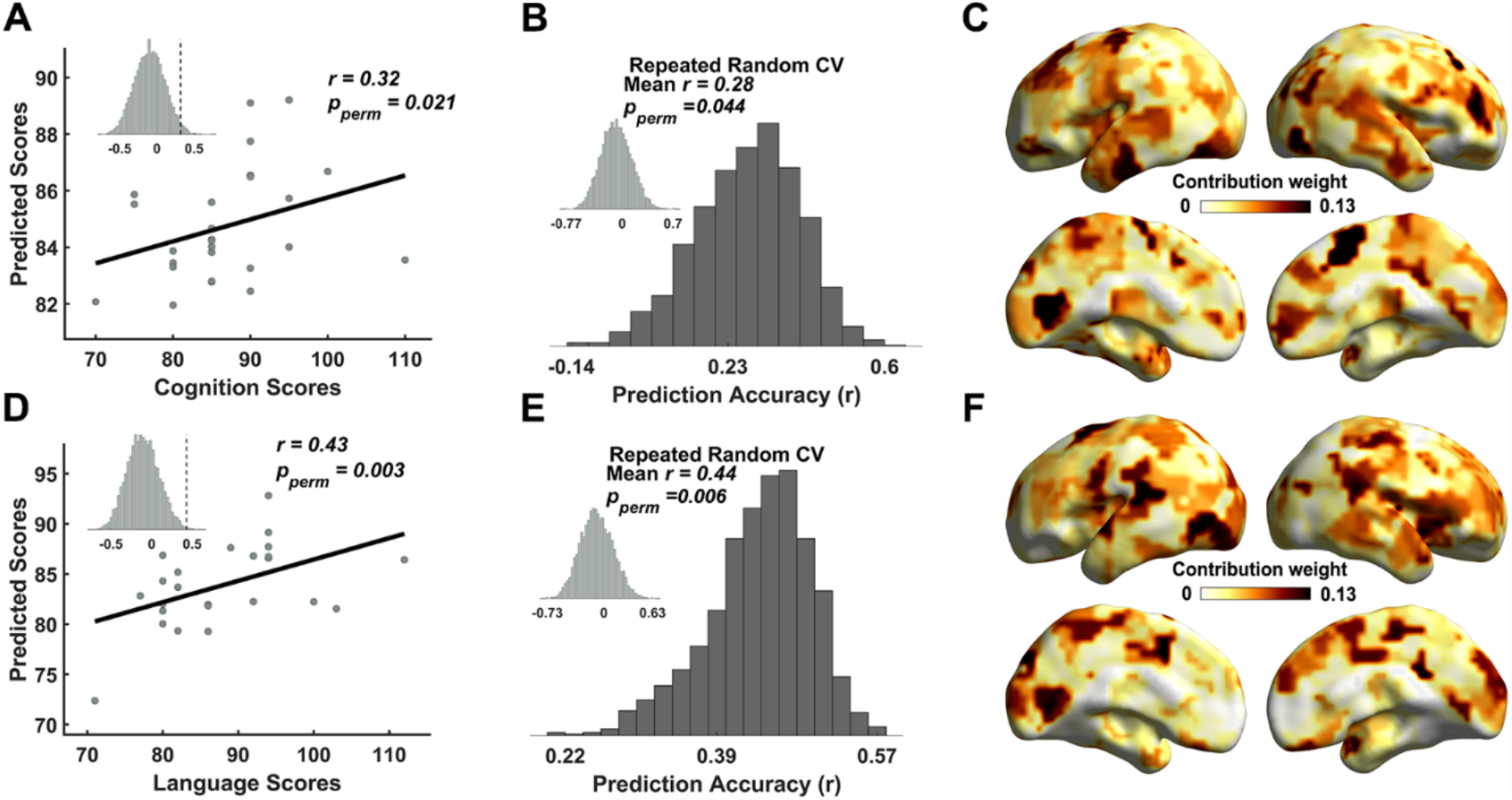
Neurocognitive outcomes prediction based on nodal modular variability at birth. (A) Prediction of individual’s Bayley cognition scores at 2 years of age. The data point represents the partial correlation between actual scores and predicted scores, corrected for effects of sex, mean FD, and time interval between birth and scan. The inset histogram shows the distribution of prediction accuracy from permutation test (*n* = 10, 000). The 10F-CV was conducted by splitting all infants into 10 subsets that were matched on age at scan. (B) Repeated random 10F-CV (1, 000 runs) demonstrated stable prediction accuracy, which was better than a null distribution with permuted data (inset histogram). (C) The absolute contribution weight of nodal regions in the cognition prediction SVR model. (D) Prediction of individual’s Bayley language scores at 2 years of age. The data point displays the partial correlation between actual scores and predicted scores, corrected for effects of sex, mean FD, and time interval between birth and scan. The inset histogram shows the distribution of prediction accuracy from permutation test (*n* = 10, 000). The 10F-CV was conducted by splitting all infants into 10 subsets that were matched on age at scan. (E) Repeated random 10F-CV (1, 000 runs) provided evidence of stable prediction accuracy, which was better than a null distribution with permuted data (inset histogram). (F) The absolute contribution weight of nodal regions in the language prediction SVR model. 10F-CV, 10-fold cross validation; SVR, support vector regression; FD, framewise displacement.

### Validation results

To assess the reliability of our results, we evaluated the effects of different analysis strategies on our main findings, including the window length (100 s), the network density (10% and 20%), and multilayer parameters (*ω* = 0.5 and 0.75; *γ* = 0.9). We found that our main findings were largely unchanged under different analysis strategies. The age-related decreases in modular variability were mainly located in the sensorimotor areas, lateral parietal cortex, and medial temporal lobe (Fig. 4, *p*_*FDR*_ < 0.05, areas delineated with yellow lines, correcting for multiple comparisons). The prediction analyses of cognitive and language scores were also significant (Table 2).

**Table 2.** Predicting neurocognitive outcomes from connectome dynamics at birth under different analysis strategies.

|  | Cognition | Language |
| --- | --- | --- |
| Window length = 100 s | $r = 0.43$, $p_{\text{perm}} = 0.007$ | $r = 0.44$, $p_{\text{perm}} = 0.005$ |
| Network density = 10% | $r = 0.29$, $p_{\text{perm}} = 0.033$ | $r = 0.34$, $p_{\text{perm}} = 0.011$ |
| Network density = 20% | $r = 0.38$, $p_{\text{perm}} = 0.012$ | $r = 0.41$, $p_{\text{perm}} = 0.003$ |
| $\omega = 0.50$, $\gamma = 1$ | $r = 0.28$, $p_{\text{perm}} = 0.029$ | $r = 0.31$, $p_{\text{perm}} = 0.016$ |
| $\omega = 0.75$, $\gamma = 1$ | $r = 0.33$, $p_{\text{perm}} = 0.021$ | $r = 0.45$, $p_{\text{perm}} = 0.002$ |
| $\omega = 1$, $\gamma = 0.90$ | $r = 0.32$, $p_{\text{perm}} = 0.022$ | $r = 0.44$, $p_{\text{perm}} = 0.002$ |

All the predictions were significant with a significance level $p_{\text{perm}} < 0.05$

**Figure 4.**
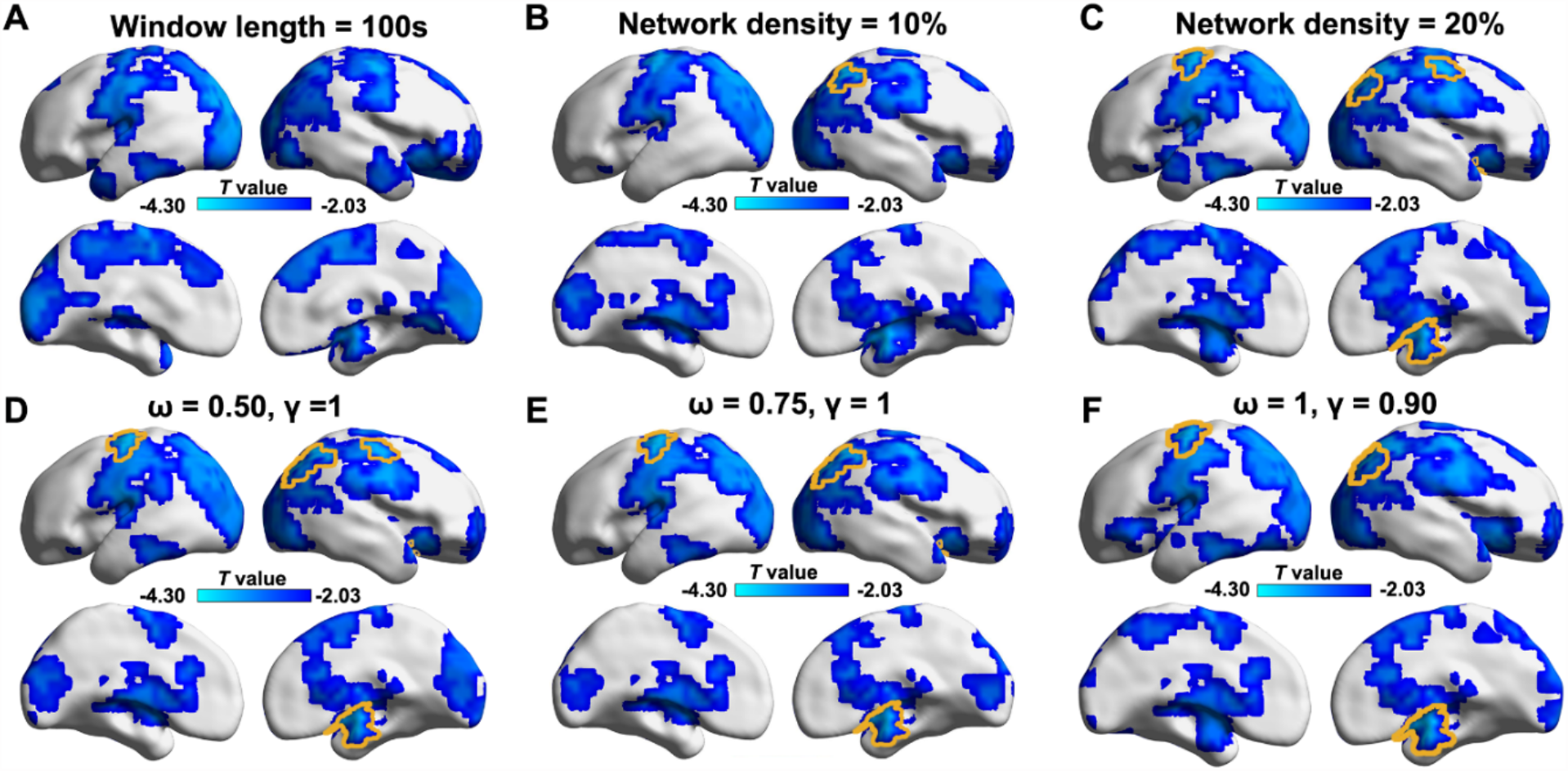
Age effects on nodal modular variability during the third trimester under different analysis strategies. (A) Window length =100 s. (B) Network density = 10%. (C) Network density = 20%. (D) Changed temporal coupling parameter (*ω*= 0.50, *γ*= 1). (E) Changed temporal coupling parameter (*ω* = 0.75, *γ*= 1). (F) Changed topological parameter (*ω* = 1, *γ*= 0.90). Here, we only display regions that show significant age-related changes (*p* < 0.05, uncorrected). Yellow curves delineate bran regions that remained significant with correction for multiple comparisons (*p*_FDR_ < 0.05). FDR, false discovery rate.

## Discussion

In this study, we revealed the emergence and maturation of brain connectome dynamics during the third trimester and their predictions on neurocognitive outcomes. Specifically, the network modular dynamics during the third trimester exhibited an adult-like spatial pattern, with higher modular switching in high-order association cortex and lower modular switching in primary regions. Moreover, the modular dynamics became progressively more stable with development in a spatially heterogeneous manner, with significant decreases mainly in the primary regions, whereas limited changes in the higher-order regions. Finally, the modular variability at birth significantly predicted the cognitive and language scores at 2 years of age. Taken together, these findings provide new insights into brain connectome dynamics during the third trimester and network mechanisms for supporting cognitive and language development in later life.

Prior studies of healthy adults and children have demonstrated the time-varying functional modular architecture during the resting state (Liao et al. 2017; Liu et al. 2020), which promotes efficient communications between networks (Zalesky et al. 2014) and a fast response to potential cognitive demands (Barbey 2018; Uddin 2021). Compared with these studies, we extended these findings to an earlier stage of life, specifically the third trimester. The heterogeneous spatial pattern of the module dynamics is similar to those observed in infants (Wen et al. 2020; Yin et al. 2020), adolescents (Lei et al. 2022), and adults (Liao et al. 2017; Liu et al. 2020). Moreover, we found that the modularity of the functional network increased with age, while the module dynamics decreased with age, which indicate the reduced dynamic communications at the system level. Similar results have been also observed in a recent study regarding module dynamics development in children and early adolescence (Lei et al. 2022). These findings, along with previous research, indicate the continuous enhancement of functional segregation during both prenatal and postnatal periods. Recent task-related studies suggest that the functional network segregation is enhanced during short-term learning to improve the task automation (Bassett et al. 2015; Finc et al. 2020). Thus, it is reasonable to assume that the gradual functional specialization of brain regions and networks during development likely underlies the neurocognitive developments (Johnson 2011; Battista et al. 2018).

Compared with the limited changes of association areas, we found that the primary areas showed higher developmental rates (i.e., significant decreases) in the connectome dynamics during the prenatal period. The divergent developmental rules across regions have also been observed for connectome dynamics in the early postnatal period (Wen et al. 2020; Yin et al. 2020). From birth to 2 years of age, the higher-order functional systems showed increased network switching with age, while the primary functional systems exhibited different developmental trends (Wen et al. 2020; Yin et al. 2020). These divergent developmental patterns across the brain may be attributed to the earlier maturation of the primary areas compared to the higher-order functional related brain regions (Cao et al. 2017a; Gao et al. 2017). Static functional network studies have revealed that the sensorimotor, visual and auditory networks showed adult-like patterns in preterm and full-term newborns, while the dorsal attention, default-mode, and frontoparietal networks are still immature at the age of 1 year and only become functionally connected in later years (Gao et al. 2015b; Gao et al. 2015a). Diffusion MRI studies have indicated that the cortical microstructure of primary sensory and motor regions matures earlier than the higher-order associative cortices (Deipolyi et al. 2005; Yu et al. 2016; Ouyang et al. 2019a; Ouyang et al. 2019b). Histological studies have also demonstrated that developmental events occur at the different timing across brain regions (Tau and Peterson 2010), wherein lamination and synaptogenesis begin earlier in the primary sensory and motor areas, and later in the prefrontal cortex (Huttenlocher 1984, 1990; Kostovic et al. 2019). All these evidences suggest the earlier development and maturation of the primary regions, which may support the basic functions for early survival in early life.

The human brain undergoes explosive growth during infancy, which is deemed to lay the critical foundation for motor, language, and cognitive development in later life (Cao et al. 2017a; Gilmore et al. 2018). Recently, a growing body of research with classical statistical methods has demonstrated the association between the intrinsic functional brain networks of infants and a broad range of cognitive abilities and behavior development later in life (Alcauter et al. 2014; Graham et al. 2016). Employing a machine learning algorithm, we found that the neurocognitive outcomes at 2 years of age can be predicted by the connectome dynamics at birth, suggesting a crucial roles of module dynamics for future developments of cognitive and language. Brain regions with large prediction contribution to cognitive scores were primarily located in the regions that have been associated with high-level cognitive functions. The lateral and medial frontal cortices are involved in the high-order cognitive processes, such as memory, attention, and decision-making (Miller and Cohen 2001). The precuneus is the core region of the default-mode network and is involved in self-awareness, episodic memory, executive functions, etc. (Buckner and DiNicola 2019). Brain regions with high prediction contribution to language scores were mainly located in the inferior frontal gyrus, supramarginal gyrus, and insular cortex. The inferior frontal gyrus and the supramarginal gyrus, known as Broca’s area and Wernicke’s area, respectively, play vital roles in language processing, such as language production, speech processing, and language comprehension (Hickok and Poeppel 2007; Hagoort 2014). Interestingly, the insular cortex showed large prediction contribution to both cognitive and language scales. The insular cortex is an integration hub that connects with extensive cortical and subcortical regions, which is primarily involved in sensory, motor control, socioemotional, language processing and cognitive functions (Oh et al. 2014; Alcauter et al. 2015; Gogolla 2017). More interestingly, these regions were also the top contributors to predicting cognitive and language outcomes in a different study from our group that used diffusion MRI-based cortical microstructure measurements as features in a largely overlapped cohort (Ouyang et al. 2020). In this study, the motor scores from Bayley-III test were not predicted, may due to the low inter-individual variability in primary motor areas associated with motor function. In our previous study, we found that the brain functional networks showed lower inter-individual variability in primary sensorimotor areas and higher variability in association regions at the third trimester (Xu et al. 2019). High inter-individual variability enables reliable prediction whereas low variability cannot capture individualized features to predict the behavior outcomes.

Several issues need to be further addressed in future research. First, preterm birth is a risk factor for the potential adverse development of the brain and behavior. Nevertheless, due to the difficulty in the fetal imaging, MRI examination of preterm infants is widely used as a substitute model to understand brain functional, structural and physiological changes during the third trimester (Cao et al. 2017b; Ouyang et al. 2017; Ouyang et al. 2019b). Some studies have shown that the effects of exposure to the environment are relatively subtle compared with the dramatic developmental changes that occur during the third trimester (Kostović 1991; Bonifacio et al. 2010), further confirming the feasibility of the substitute model. In this study, we have also used a general linear model to regress the time duration between birth and the scan to eliminate the ex-utero environmental influences. Our findings need further confirmation when high-quality fetal imaging studies are available, which can provide a more direct brain developmental curves in utero (van den Heuvel and Thomason 2016). Second, different sleep states among infants during the rs-fMRI scanning may introduce bias into our findings. In this study, all the infants were well-fed and imaged as soon as possible after falling asleep, which would minimize the differences in sleep states across infants. Third, our study has a small sample size because scanning neonates without sedation is very challenging. We believe that with the data release of the Baby Connectome Project (Howell et al. 2019) and the developing Human Connectome Project (Fitzgibbon et al. 2020), our prediction model could be further verified with these new infant neuroimaging datasets (Scheinost et al. 2022). Finally, previous studies on adults have shown that the dynamic functional connectivity is structurally constrained by white matter tracts (Liao et al. 2015; Zhang et al. 2016). The development of static functional connectivity is tightly coupled with regional cerebral blood flow, which delivers nutrients to different brain regions (Liang et al. 2013; Yu et al. 2023).However, how anatomical substrates (e.g., cortical morphology and white-matter structural connectivity) and brain blood supply contribute to the development of the connectome dynamics warrants further investigation.

## Acknowledgements

We would like to thank Drs. Miao Cao and Jiaying Zhang for their insightful comments on the manuscript. This work was supported by Science, Technology and Innovation (STI) 2030-Major Projects (2021ZD0201700), the Natural Science Foundation of China (Grant Nos. 31830034, 82102131, 81971690, 82021004, 82202245), and the Tang Scholar Award of Beijing Normal University, and National Institute of Health (Grant Nos. R01MH092535, R01MH125333, R01EB031284, R01MH129981, R21MH123930, and P50HD105354).

## Notes

### Competing Interest Statement

The authors have declared no competing interest.

### Summary of Updates

authorship updated

